# State-dependent, frequency-specific modulation of electrocortical traveling waves by anesthesia

**DOI:** 10.64898/2026.08.11.744271

**Authors:** Duan Li, Anthony G. Hudetz

**Author notes:** Corresponding Author: Duan Li, Ph.D., Center for Consciousness Science, Department of Anesthesiology University of Michigan Medical School, North Campus Research Complex (NCRC), Building 16 2800 Plymouth Road, Ann Arbor, MI 48109.

## Abstract

Emerging evidence suggests that cortical activity is organized in traveling waves that coordinate neural activity across space and time. How anesthesia alters these waves remains underexplored. We recently showed that cortical activity undergoes spontaneous state transitions at steady-state anesthetic concentrations including a paradoxical state exhibiting awake-like spectral properties during deep anesthesia. Here, we investigated traveling wave dynamics across spontaneous cortical states using hemispheric electrocorticography in rats anesthetized with desflurane at inhaled concentrations of 6, 4, 2, and 0%. Compared with the awake state, delta-band traveling waves in cortical states predominantly associated with 4-6% desflurane were more frequent and exhibited more stereotyped propagation patterns, characterized by a greater prevalence of planar waves and a corresponding reduction in source/sink wave patterns. The occurrence rate and pattern complexity of theta- and gamma-band waves remained largely unchanged, whereas the propagation direction of planar waves became more variable. Feedforward-feedback organization was also altered: compared with the awake state, the feedback-dominance of theta-band diminished, and the feed-forward dominance of gamma-band was attenuated. Despite occurring predominantly in deep anesthesia associated with behavioral unresponsiveness, traveling-wave dynamics of the paradoxical state exhibited partial, frequency-dependent shifts toward those observed in the awake state. These findings demonstrate that spontaneous cortical states under anesthesia are associated with frequency-dependent reorganization of cortical traveling waves and identify the paradoxical state as a distinct dynamical regime of deep anesthesia.

**Significance Statement:** Anesthesia is commonly thought to alter cortical dynamics progressively with increasing anesthetic depth, yet cortical activity can transition spontaneously between distinct states even at constant anesthetic concentrations. Here, we show that cortical states spectrally derived from the electrocorticogram of rats are associated with distinct frequency-specific organization of cortical traveling waves, revealing spatiotemporal dynamics beyond conventional spectral measures. Notably, a paradoxical state, occurred predominantly in deep anesthesia associated with behavioral unresponsiveness, exhibited traveling-wave dynamics that approached those observed during wakefulness. These findings demonstrate that cortical traveling-wave organization changes dynamically with brain state rather than anesthetic concentration alone. They suggest that structured cortical dynamics can emerge during deep anesthesia, providing new insights into large-scale cortical dynamics associated with anesthetic modulation of consciousness.

## Introduction

Neuronal oscillations are fundamental features of brain activity and are closely associated with distinct behavioral and physiological states. Traditionally, these oscillations have been studied as markers of synchrony or functional connectivity by quantifying statistical dependencies between recording sites. However, growing evidence suggests that cortical activity often manifests as traveling waves—spatiotemporally organized patterns of neural activity that propagate across the cortex in a coordinated manner (Muller et al., 2018). Traveling waves have been observed across spatial scales ranging from local cortical circuits to large-scale brain networks, as well as across species, behavioral states, and both spontaneous and stimulus-evoked activity. The mechanisms underlying these waves likely vary across spatial scales and are thought to involve local circuitry, anatomical geometry, large-scale connectivity, and intrinsic neural dynamics (Cruddas et al., 2026; Muller et al., 2026). Traveling waves have been implicated in sensory processing (Davis et al., 2020; Aggarwal et al., 2022; Aggarwal et al., 2024), motor coordination (Balasubramanian et al., 2023; Liang et al., 2023a), and memory consolidation during sleep (Muller et al., 2016; Xu et al., 2025), highlighting their functional relevance across diverse brain states and processes.

Understanding how general anesthesia alters cortical dynamics offers a promising approach for uncovering neural mechanisms underlying consciousness. General anesthesia profoundly reshapes cortical activity in a frequency-dependent manner, often enhancing low-frequency activity and reorganizing higher-frequency dynamics and their coordination across cortical regions (Purdon et al., 2015; Hudetz and Mashour, 2016). Recent studies have examined traveling wave dynamics during general anesthesia. However, this work has primarily focused on low-frequency activity, including slow oscillations measured with extracellular local field potential recordings (Dasilva et al., 2021; Pazienti et al., 2022; Pazienti et al., 2025) and delta-band waves measured with hemisphere-scale optical voltage imaging (Townsend et al., 2015; Liang et al., 2021; Liang et al., 2023b). A number of studies have examined traveling waves at higher frequencies (e.g. 8-30 Hz) or across broadband activity, but typically within spatially restricted cortical regions (Bhattacharya et al., 2022; Bardon et al., 2025; Zarr et al., 2026). Thus, how anesthesia modulates large-scale cortical traveling waves across distinct frequency bands remains poorly understood.

Although general anesthetics alter cortical activity in a dose-dependent manner, local brain states can fluctuate dynamically even at constant anesthetic drug levels. Several studies have shown that cortical activity during general anesthesia spontaneously transitions among multiple discrete brain states that do not show a one-to-one correspondence with anesthetic concentration (Hudson et al., 2014; Li et al., 2019; Lee et al., 2020; Li and Hudetz, 2025, 2026). In particular, one state occurring predominantly during deep anesthesia exhibits unexpectedly low delta activity and high spatiotemporal complexity (Li and Hudetz, 2026). This so-called paradoxical state raises the intriguing possibility that neural dynamics at a high anesthetic concentration typically associated with behavioral unresponsiveness (Imas et al., 2005b) can partially shift toward those observed during wakefulness. Beyond measures of signal complexity, traveling waves provide information about how cortical activity is spatially organized and propagates across the cortex. Examining their dynamics across discrete cortical states may therefore provide further insight into state-dependent cortical reorganization under anesthesia, which remains incompletely understood.

To fill these knowledge gaps, we investigated how traveling wave dynamics in hemispheric electrocorticogram (ECoG) were altered by the general anesthetic desflurane, administered at inhaled concentrations of 6%, 4%, 2%, and 0% in rats. Specifically, we examined changes in wave occurrence, wave pattern diversity, wave types (planar, rotating, and source/sink), and planar wave directional organization in the delta, theta, and gamma bands across discrete cortical states, including the paradoxical state. We found that desflurane anesthesia induced frequency-and state-dependent reorganization of traveling wave dynamics. Despite occurring predominantly at the highest anesthetic concentration studied here, the paradoxical state showed partial reversal of spatiotemporal cortical organization toward wakefulness, distinguishing it from canonical deep anesthetic state and extending its previous characterization based on spectral power and complexity (Li and Hudetz, 2026).

## Materials and methods

The study was approved by the Institutional Animal Care and Use Committee of the University of Michigan. All procedures were performed in accordance with the *Guide for the Care and Use of Laboratory Animals* (National Research Council, 2011) and reported in accordance with the *Animal Research: Reporting of In Vivo Experiments* guidelines (Percie du Sert et al., 2020).

### Experimental design

We analyzed electrocorticogram (ECoG) data obtained in a previous study (Li and Hudetz, 2026). Briefly, eight adult male Sprague-Dawley rats (7-9 weeks old, 300-350 g) were equipped with chronically implanted 32-channel flexible polymer electrode arrays arranged in a 4×8 grid (300-µm site diameter, 1-mm inter-site spacing; Neuromicrosystems, Budapest, Hungary) (Fedor et al., 2020). The array was positioned with its long axis parallel to the midline and primarily covered the primary and secondary somatosensory cortex and adjacent parietal association areas (**Figure 1A**). The reference electrode was placed in the nasal sinus, and a stainless-steel screw over the cerebellum was placed in the cranium for ground. Each rat underwent two to three experiments to minimize animal use. Five experiments were excluded due to excessive electrical noise (n=4) or a very low incidence of traveling wave episodes (n=1), leaving 13 experiments (two from each of five rats, and one from each of three rats). Animals were placed in a ventilated chamber allowing free movement. Desflurane was administered at stepwise decreasing concentrations (6%, 4%, 2%, and 0%) in 30% O_2_ balanced with N_2_ (**Figure 1B**), with continuous monitoring of anesthetic concentration (POET IQ2, Criticare Systems, Inc., Waukesha, WI, USA). Although behavioral responsiveness was not tested in the present study, our previous work showed that loss of righting reflex occurred at desflurane concentrations between 3.5% and 5.0% (Imas et al., 2005b). Loss of righting reflex is commonly used as a behavioral surrogate for loss of consciousness in rodents, analogous to the loss of response to verbal commands used to assess loss of consciousness in humans (Franks, 2008). Accordingly, 2% desflurane was presumed to represent sedation with preserved consciousness, 4% to fall within the transition range between conscious and unconscious states, and 6% to represent unconsciousness. Core body temperature was maintained at 37 °C using subfloor heating. Each anesthetic concentration was maintained for 75 minutes.

**Figure 1.**
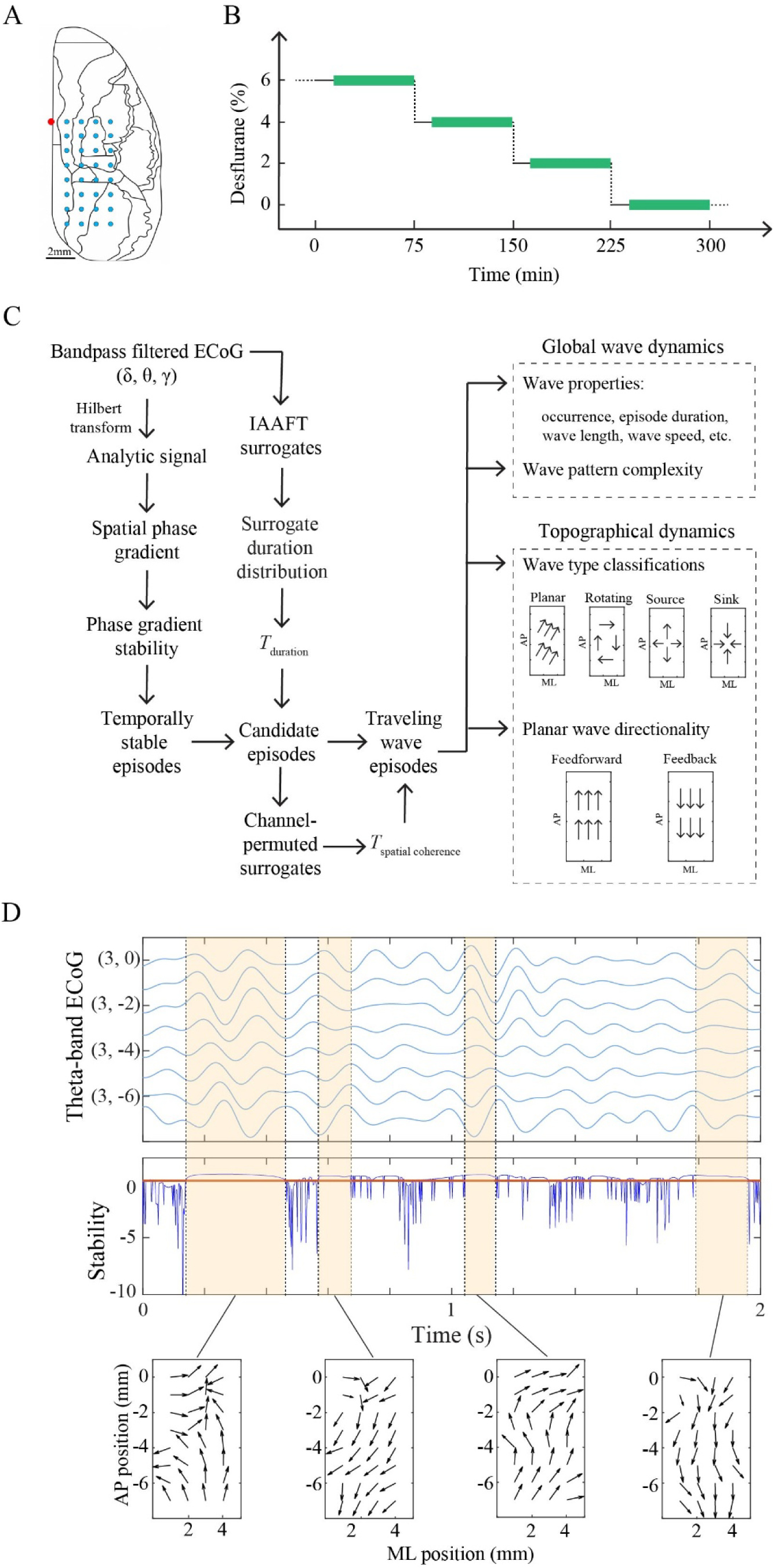
Schematic overview of the experimental design and analysis pipeline. **A**. Placement of the high-density ECoG grid (8×4 electrodes) implanted over the right hemisphere of the rat cortex. The red dot indicates bregma. **B**. Desflurane was administered in stepwise reductions of steady-state concentrations. After a 15-min equilibration period at each concentration, spontaneous ECoG activity was recorded for 1 h (green bars). **C**. Analysis pipeline for detecting and characterizing traveling wave episodes. Instantaneous phase was obtained from bandpass-filtered signals in the delta (0-4 Hz), theta (4-8 Hz), and gamma (25-55 Hz) bands using the Hilbert transform. Spatial phase gradients were then computed to characterize wave propagation. The temporal stability of these phase-gradient patterns was quantified based on the similarity of phase gradient vectors between consecutive time points. Candidate episodes were identified by thresholding temporal stability and retaining episodes whose durations exceeded an IAAFT surrogate-derived threshold (*T*_duration_). Spatial coherence was then evaluated based on the spatial variability of the phase-gradient pattern, and episodes with spatial variability below the corresponding channel-permuted surrogate-derived threshold (*T*_spatial coherence_) were retained as traveling wave episodes. For each episode, occurrence rate, wave properties (duration, wavelength, and propagation speed), wave pattern complexity, and wave type (planar, rotating, source/sink) were quantified. **D**. Representative example of traveling wave episodes. Top: theta-band ECoG traces from one column of electrodes along the anterior-posterior axis. Middle: temporal profile of phase gradient stability, with shaded regions indicating candidate episodes exceeding the mean stability (red line) and IAAFT surrogate-derived duration threshold. Bottom: representative phase gradient patterns for each episode, illustrating propagation direction (arrow orientation) and temporal consistency (arrow length). AP, anteroposterior, ML, mediolateral. IAAFT, iterated amplitude-adjusted Fourier transform.

### Electrophysiological signal preprocessing

Spontaneous ECoG signals were recorded for 60 minutes following a 15-min equilibration period at each anesthetic concentration. Signals were sampled at 1 kHz and exported to MATLAB (version 2024b; MathWorks, Inc., Natick, MA). Preprocessing followed our prior work (Li and Hudetz, 2026). Briefly, signals were detrended, low pass filtered at 55 Hz, visually inspected for poor-quality channels (which were interpolated), and re-referenced to the common average. Artifacts were identified using an amplitude threshold and removed (±1 s around events). The number of interpolated channels (median [IQR]) was 1[0, 2] at 6%, 4%, and 2% desflurane, and 1[1, 2] at 0% desflurane. The percentage of rejected data was 0 [0, 0] %, 0 [0, 0.29] %, 2.43 [1.66, 5.05] %, 3.70 [3.38, 5.66] % at 6%, 4%, 2%, and 0% desflurane, respectively. The remaining data were concatenated for further analysis.

### Definition of cortical states

Cortical states were defined based on our previous study, in which spontaneous ECoG activity was clustered using power spectrograms (Li and Hudetz, 2026). Briefly, time-resolved power spectrograms were computed using 2-s windows with 1-s overlap and smoothed over 30 consecutive windows, then reduced using principal component analysis, followed by density-based clustering to identify discrete activity patterns within each experiment. Clustering was performed on concatenated data from different anesthetic conditions to allow delineation of different brain states independent of the actual anesthetic level. The clusters were then aligned across experiments and mapped to a common set of cortical states based on their spectral profiles and occupancy across anesthetic concentrations. The resulting cortical states exhibited distinct spectral profiles across the power spectrum, including the delta, theta and gamma bands, and were ordered according to the progression in delta power across cortical states. Subsequent traveling-wave analyses were performed separately within the delta, theta, and gamma bands.

### Detection of traveling waves

In this study, traveling waves were defined as temporally stable and spatially coherent phase gradient patterns across the electrode array, reflecting propagating oscillatory activity (**Figure 1C**). ECoG signals were bandpass filtered into delta (0-4 Hz), theta (4-8 Hz), and gamma (25-55 Hz) bands using a 3^rd^-order Butterworth filter implemented with a zero-phase forward and reverse algorithm (*filtfilt.m* function in Matlab). The analytic signal was then obtained using the Hilbert transform. Spatial phase gradients were computed at each time point using complex multiplication between adjacent electrodes (Muller et al., 2016), using the wave toolbox (https://github.com/mullerlab/wave-matlab). Specifically, the analytic signal at each electrode was multiplied by the complex conjugate of its neighboring electrode, and this operation was applied iteratively along the anterior-posterior and medial-lateral directions of the grid. The angle and magnitude of the resulting vector represent the direction and strength of local phase propagation, respectively.

To identify temporally stable traveling wave patterns, we quantified the temporal stability of phase gradient vectors between consecutive time points (Ito et al., 2007; Das et al., 2026). Stability was defined as the negative mean absolute difference in phase gradient vectors across electrodes between successive time points (Das et al., 2026):

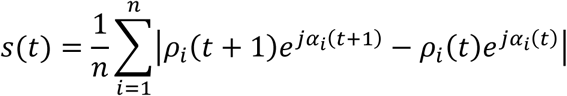

where *ρ_i_* and *α_i_* denote the magnitude and angle of the phase gradient at electrode *i*, respectively, and *n* is the number of electrodes. Stability values were z-scored, and candidate wave episodes were defined as periods during which stability exceeded zero.

To establish statistical thresholds for wave episode duration and spatial coherence, surrogate data were generated using the iterative amplitude-adjusted Fourier transform (IAAFT) algorithm, applied independently to each channel. Surrogates were constructed within non-overlapping 10-s windows, preserving channel-wise amplitude distributions and power spectra while disrupting phase relationships. For each experiment, the durations of surrogate wave episodes were pooled across windows to form a null distribution, and duration thresholds were defined as the 99^th^ percentile of this distribution (**Figure 1D** and **Figure S1**). Spatial coherence was then quantified from the spatial variability of the phase-gradient pattern, defined as the spatial gradient of the phase-gradient vectors across the electrode array. Spatial coherence thresholds were defined as the 99^th^ percentile of the corresponding null distributions obtained from channel-shuffled surrogate data. Finally, only candidate episodes that exceeded the duration threshold and exhibited spatial variability below the corresponding spatial coherence threshold were retained as traveling waves (**Figure 1D**). For each episode, a representative phase gradient pattern was obtained by circularly averaging phase gradient directions and separately averaging phase gradient magnitudes over all time points within the episode.

### Quantification of wave properties

For each frequency band, experiment, and cortical state, we quantified the occurrence rate, episode duration, wavelength, and propagation speed of traveling waves. The occurrence rate was defined as the proportion of time occupied by wave episodes relative to the total recording duration within each state. Episode duration was defined as the temporal length of each wave episode and averaged across episodes within each experiment and cortical state.

For each traveling wave episode, wavelength at each site was calculated as the reciprocal of the mean phase gradient magnitude: 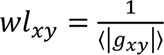, where 〈·〉 denotes averaging over time points within the episode, and *x* and *y* indicate locations in the mediolateral and anterior-posterior directions, respectively. Wave speed was computed as: 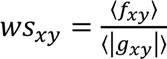, where *f_xy_* is the instantaneous temporal frequency at each time point, estimated as the temporal derivative of phase, 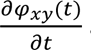and then averaged over time within the episode.

### Quantification of wave pattern complexity

The diversity of spatial wave patterns across episodes was quantified for each experiment, cortical state, and frequency band using wave pattern complexity derived from the episode-by-episode similarity matrix. Similarity between episodes was measured using Earth Mover’s Distance (EMD), which quantifies the minimal cost of transforming one spatial distribution into another (Rubner et al., 2000; Aggarwal et al., 2024). Pairwise EMD was computed using the Matlab implementation *emd.m* (Yilmaz, 2009).

Each traveling wave episode was represented by the spatial distribution of phase gradient directions across the electrode array. At each electrode location (*x*, *y*), the phase gradient direction *θ_xy_* was encoded as a four-dimensional feature vector [cos(*θ_xy_*), sin(*θ_xy_*), *x*, *y*], representing both propagation direction and spatial organization of the wave pattern. To reduce computational complexity, feature vectors were clustered using *k*-means (k = 5, cityblock distance, 100 replicates) to generate compact signatures composed of weighted clusters

{*s_j_* = (*m_j_*, *w_j_*)}, where *m_j_* denotes the cluster centroid, *w_j_* the proportion of electrodes assigned to that cluster, and 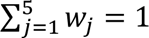. The EMD between two episodes was computed as the optimal transport cost between their corresponding signatures.

The ground distance was defined using the cityblock metric:

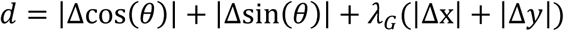

where Δ denotes the difference between corresponding feature components. The weighting factor *λ_G_* balances the contribution of directional and spatial components and was defined as 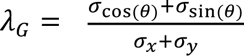, where *σ* denotes the standard deviation of each feature computed across all episodes. The resulting pairwise distance matrix was converted to a similarity matrix by subtracting each distance from the maximum matrix value, such that higher values indicated greater similarity between episodes. The similarity matrix was then normalized using min-max scaling separately for each experiment and cortical state to preserve the relative organization of wave patterns while minimizing differences in absolute distance across cortical states.

The similarity matrix was decomposed using singular value decomposition. The normalized singular value spectrum was used to characterize the dimensionality of the wave repertoire. The largest singular value reflects the dominant mode of the wave repertoire, whereas the remaining singular values capture additional independent modes contributing to repertoire diversity.

Shannon entropy of the normalized singular values was computed as the primary measure of wave pattern complexity, with higher entropy indicating a richer and more diverse repertoire of traveling wave patterns. As complementary measures, we also computed the percentage of singular values required to explain 90% of the variance and the participation ratio, both estimating the effective dimensionality of the wave repertoire.

To account for variability in the number of episodes across experiments and states, and to limit computational burden in states with large numbers of episodes, we controlled sample size by subsampling a fixed number of episodes. The target sample size was defined as the minimum across cortical states of the median number of episodes per experiment. For each experiment and state, if the number of available episodes exceeded this value, a random subset of episodes was selected; otherwise, all episodes were used. To reduce sampling variability, wave pattern complexity measures were computed over 12 independent realizations and averaged. This procedure was performed separately for each frequency band.

### Classification of wave types

To characterize the spatial organization of traveling waves, each episode was classified into canonical wave types based on the topology of its phase gradient field.

Classification of planar waves required two criteria. First, the distribution of phase gradient directions across electrodes was tested using the Rayleigh test for circular uniformity (Aggarwal et al., 2024), implemented in the Circular Statistics Toolbox (Berens, 2009), and required to be significant (*p* < 0.05). Second, the spatial consistency of propagation directions, quantified by the mean resultant vector length (1 - circular variance), was required to exceed the 99^th^ percentile of an experiment-specific surrogate distribution. The surrogate distribution was generated by randomly shuffling channel locations of ECoG signals within each episode and pooling the resulting values across episodes.

Rotating waves and source/sink waves were identified following (Muller et al., 2016). Rotating waves were detected from the curl of the phase gradient field by identifying a putative rotation center and computing the circular-circular correlation between signal phase and rotation angle.

Source and sink patterns were detected from the divergence of the phase gradient field by identifying locations of maximal positive (source) or negative (sink) divergence and computing the circular-linear correlation between signal phase and radial distance from the identified center. Statistical significance was assessed relative to a channel-shuffled surrogate distribution as described for planar waves.

Because wave types are not mutually exclusive, classification was performed sequentially as planar, rotating, and source/sink types. Episodes meeting none of these criteria were classified as atypical waves. For each experiment, cortical state, and frequency band, the proportion of episodes belonging to each wave type was quantified, providing a complementary description of state-dependent changes in wave organization.

### Characterization of planar wave organization

For each planar wave episode, the mean propagation direction was obtained by circularly averaging phase gradient directions across electrodes. Across cortical states and frequency bands, the distribution of mean propagation directions exhibited a consistent bimodal structure.

Accordingly, planar waves were classified as feedforward (FF, posterior-to-anterior) when the mean direction fell within (0, π) and feedback (FB, anterior-to-posterior) when it fell within (π, 2π).

For each experiment, cortical state, and frequency band, we quantified the preferred propagation direction (defined as the circular average of the mean propagation directions across episodes), directional consistency (defined as the mean resultant vector length of the mean propagation directions across episodes), and occurrence rate of FF and FB waves separately. Directional asymmetry was quantified by comparing the occurrence rates and directional consistency of FF and FB waves. As supplementary analyses, we also quantified episode frequency (number of episodes per second), mean episode duration, mean wavelength, and mean propagation speed separately for FF and FB waves, together with the FB/FF ratios for each measure.

### Statistical analysis

Linear mixed-effects models were fitted in IBM SPSS Statistics version 30.0 for Windows (IBM Corp., Armonk, NY) to assess state-dependent traveling wave properties across experiments, with cortical state as a fixed effect and a random intercept for each experiment to account for inter-experiment variability. All models were fitted using restricted maximum likelihood estimation. Post hoc pairwise comparisons were performed using two-tailed paired Student’s *t*-tests with Bonferroni correction. Comparisons were restricted to two prespecified contrast families: each state versus State 1 and each state versus State 7 (six comparisons per family). For each comparison, estimated mean difference, 95% confidence interval, and associated *p*-values were reported. Directional asymmetry between FF and FB waves was additionally assessed using Wilcoxon signed-rank tests to determine whether the FB/FF ratio for each wave property differed from 1, with Bonferroni correction applied across the seven cortical states. All other analyses were performed using MATLAB, unless otherwise specified. The normality of the data was assessed using the Lilliefors-corrected Kolmogorov-Smirnov test, and a *p* value < 0.05 was considered statistically significant.

## Results

### Cortical activity organizes into discrete, recurrent states across anesthetic levels

Anesthetic concentration is commonly used as a proxy for brain state, yet cortical activity exhibits substantial variability even at stable concentrations. We therefore examined cortical activity using a data-driven state classification approach as previously reported (Li and Hudetz, 2026). A representative experiment illustrates the systematic changes in spectral power during the stepwise reduction of desflurane concentration (**Figure 2A**). Clustering power spectrograms across all anesthetic concentrations identified seven cortical states with distinct spectral properties that were consistently observed across experiments. Across States 1-5, delta power progressively increased while theta and gamma power decreased, broadly corresponding to anesthetic depth from awake (State 1), to light anesthesia (States 2-3), intermediate anesthesia (State 4), and deep anesthesia (State 5) (**Figure 2B-D**). However, these states were not uniquely associated with specific anesthetic concentrations (see **Table S1** for a summary of cortical states across all experiments). State 6 corresponded to burst suppression and, together with State 5, usually occurred at the deepest level of anesthesia studied here (**Figure 2B, C**). Despite also occurring predominantly at this anesthetic level, State 7 was a paradoxical state, exhibiting reduced delta power (*p*=0.001 vs. State 5) and elevated theta (*p*=0.023 vs. State 1; not significant after Bonferroni correction) and gamma power (*p*=0.004 vs. State 5) (**Figure 2B, D**). These states were then used to examine the effects of anesthesia on traveling wave dynamics by first comparing states broadly associated with different anesthetic depths (States 1-6, hereafter referred to as canonical states; see also **Figure S2** for concentration-dependent modulation of traveling wave dynamics) and then comparing the paradoxical State 7 with these canonical states.

**Figure 2.**
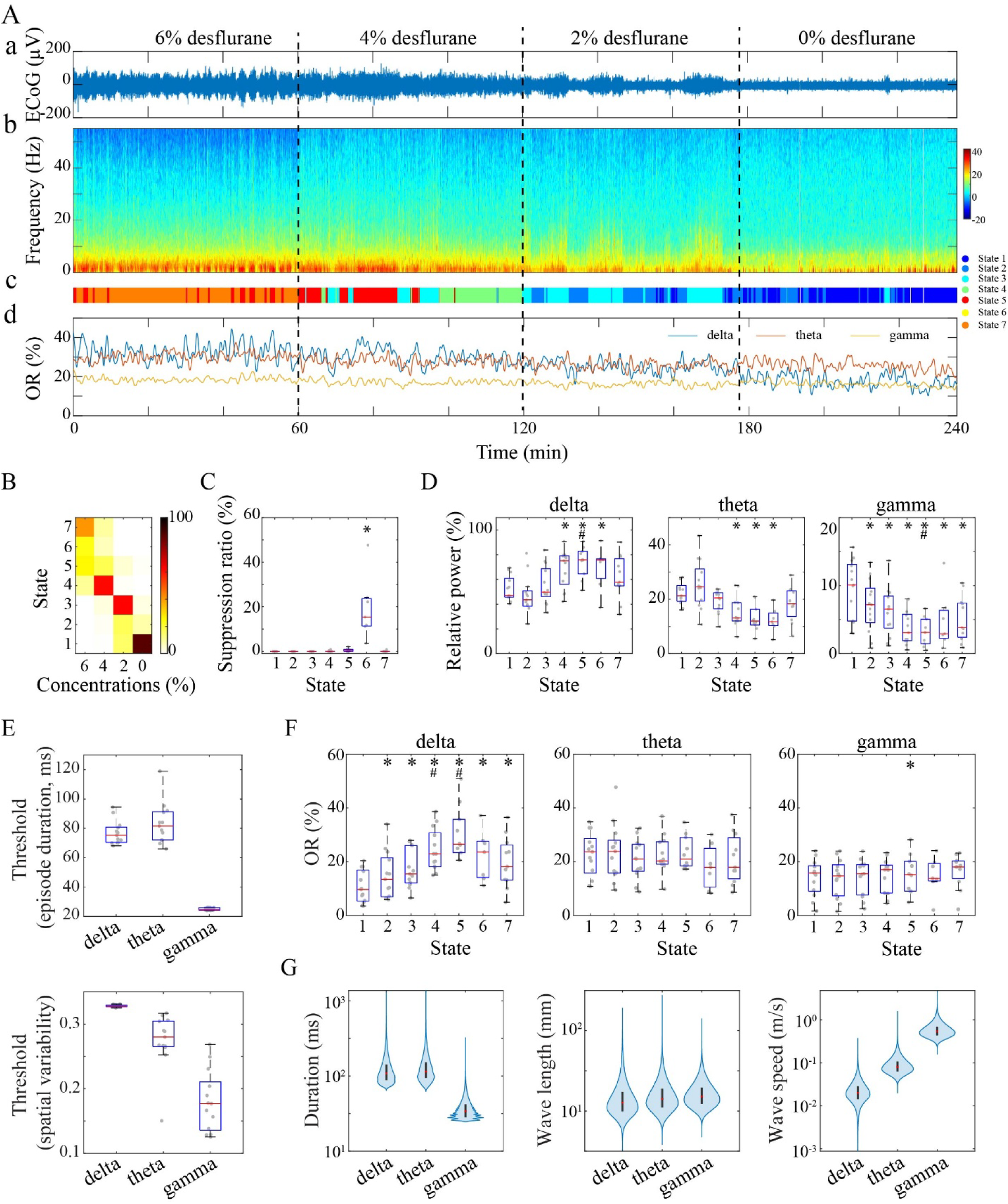
Classification of cortical states and state-dependent occurrence of traveling waves. **A**. Identification of seven cortical states in a representative experiment: (a) example ECoG signal from an occipital channel, (b) average power spectrogram across recording channels, (c) corresponding color-coded state labels, (d) time course of traveling wave occurrence rate (OR) in the delta, theta and gamma bands, computed using a 30-s sliding windows with 1-s steps. **B**. State occurrence rates across desflurane concentrations. **C**. Suppression ratio across states. **D**. State-dependent changes in band-limited power in the delta, theta and gamma bands. **E**. Frequency-specific statistical thresholds for traveling wave episode duration and spatial coherence derived from surrogate data. **F**. Occurrence rate of traveling wave episodes across cortical states for each frequency band. **G**. Wave properties of traveling wave episodes, including episode duration, wavelength, and propagation speed. In C-F, boxplots indicate the median and interquartile range (IQR); whiskers extend to the most extreme values, and individual experiments are shown as dots. In G, violin plots show the distribution of wave properties across episodes; red dots indicate the median and black bars indicate the IQR. * *p*<0.05/6 vs. State 1. # *p*<0.05/6 vs. State 7, linear mixed-effects models.

### Delta-band wave occurrence is state-dependent

Traveling waves in spontaneous ECoG activity were frequently observed as spatially coherent propagation patterns that persisted for tens to hundreds of milliseconds and recurred over time (**Figure 1D**). In contrast, waves expected by chance were typically short-lived and exhibited less coherent phase-gradient organization (**Figure S1**). To define reliable traveling-wave episodes for subsequent analyses, we focused on episodes exhibiting temporally stable phase-gradient patterns, with durations exceeding those expected from phase-shuffled data, and spatial variability in the phase-gradient pattern lower than that expected from channel-shuffled data (i.e., greater spatial coherence). The surrogate-derived detection thresholds differed across frequency bands. Compared with gamma-band waves, lower-frequency waves required longer durations but were permitted greater spatial variability to be identified as traveling waves. The median duration thresholds were 75 ms (delta), 82 ms (theta), and 25 ms (gamma), whereas the corresponding spatial-variability thresholds were 0.33, 0.28, and 0.18, respectively (**Figure 2E**).

In a representative experiment, delta-band wave occurrence closely tracked cortical state transitions, whereas theta- and gamma-band wave occurrence remained relatively stable (**Figure 2Ad**). Across experiments, delta-band wave occurrence followed a state-dependent pattern similar to that of delta power, was higher in all anesthetic states than in State 1 (States 2-6 vs. State 1, all *p*≤0.007) and reached its highest level in State 5. This trend was partially reversed in State 7, where delta-wave occurrence remained higher than in State 1 (8.31 [5.30, 11.73], *p*<0.001) but was significantly lower than in State 4 (−5.09 [−8.30, −1.89], *p*=0.002) and State 5 (−10.82 [−14.29, −7.34], *p*<0.001). In contrast, theta- and gamma-band wave occurrence showed little or no modulation across states (theta: *p*=0.107; gamma: *p*=0.018) (**Figure 2F, Table S2**), despite marked state-dependent changes in band-limited power. In addition, we examined the lag-dependent temporal overlap between traveling waves in different bands. Temporal overlap was generally weak, showing only a modest increase during deep anesthesia that was absent in State 7 (**Figure S3**).

The state-dependent increase in delta-band wave occurrence resulted from increases in both episode frequency (i.e. number of episodes per second) and episode duration (**Figure S4**). Beyond these state-dependent differences, wave duration varied more markedly across frequency bands. Consistent with the frequency-dependent duration thresholds (**Figure 2E**), delta and theta waves exhibited similarly long durations (delta: 108 [88, 142] ms; theta: 114 [94, 152] ms), whereas gamma waves exhibited shorter durations (34 [28, 42] ms) (**Figure 2G**). In contrast, the spatial scale of traveling waves was largely conserved across frequency bands, with comparable wavelengths (delta: 12.66 [9.92, 17.30] mm; theta: 14.13 [11.13, 18.94] mm; gamma: 15.15 [12.23, 19.51] mm). Consequently, propagation speed increased with temporal frequency, from 0.02 [0.01, 0.03] m/s in the delta band to 0.08 [0.06, 0.11] m/s in the theta band and 0.54 [0.44, 0.70] m/s in the gamma band (**Figure 2G**).

Together, these results indicate that i) traveling waves are ubiquitous in spontaneous cortical activity, ii) the occurrence of delta-band traveling waves is selectively modulated by cortical state, and iii) this state-dependent increase is partially reversed in the paradoxical state despite occurring during deep anesthesia.

### Delta-band wave patterns are more stereotyped and planar under anesthesia

To examine how cortical state influences the organization of traveling waves, we analyzed the repertoire of traveling wave episodes (i.e., individual temporally continuous, spatially coherent wave events) across cortical states using two complementary approaches: (1) characterizing the structure of the wave-pattern space based on between-episode similarity, and (2) quantifying the relative proportions of different wave types within each cortical state.

Low-dimensional embedding of delta-wave patterns based on between-episode similarity revealed a broad distribution of wave patterns in the awake state (State 1), consistent with a diverse wave repertoire. In contrast, wave patterns were more constrained in the deep anesthetic state (State 5), indicating a more stereotyped organization (**Figure 3A**). Consistent with these observations, singular value analysis of the similarity matrix revealed greater dominance of the leading singular value together with lower singular value entropy (wave-pattern complexity), indicating that the wave repertoire was more stereotyped and less diverse (**Figure 3B**).

**Figure 3.**
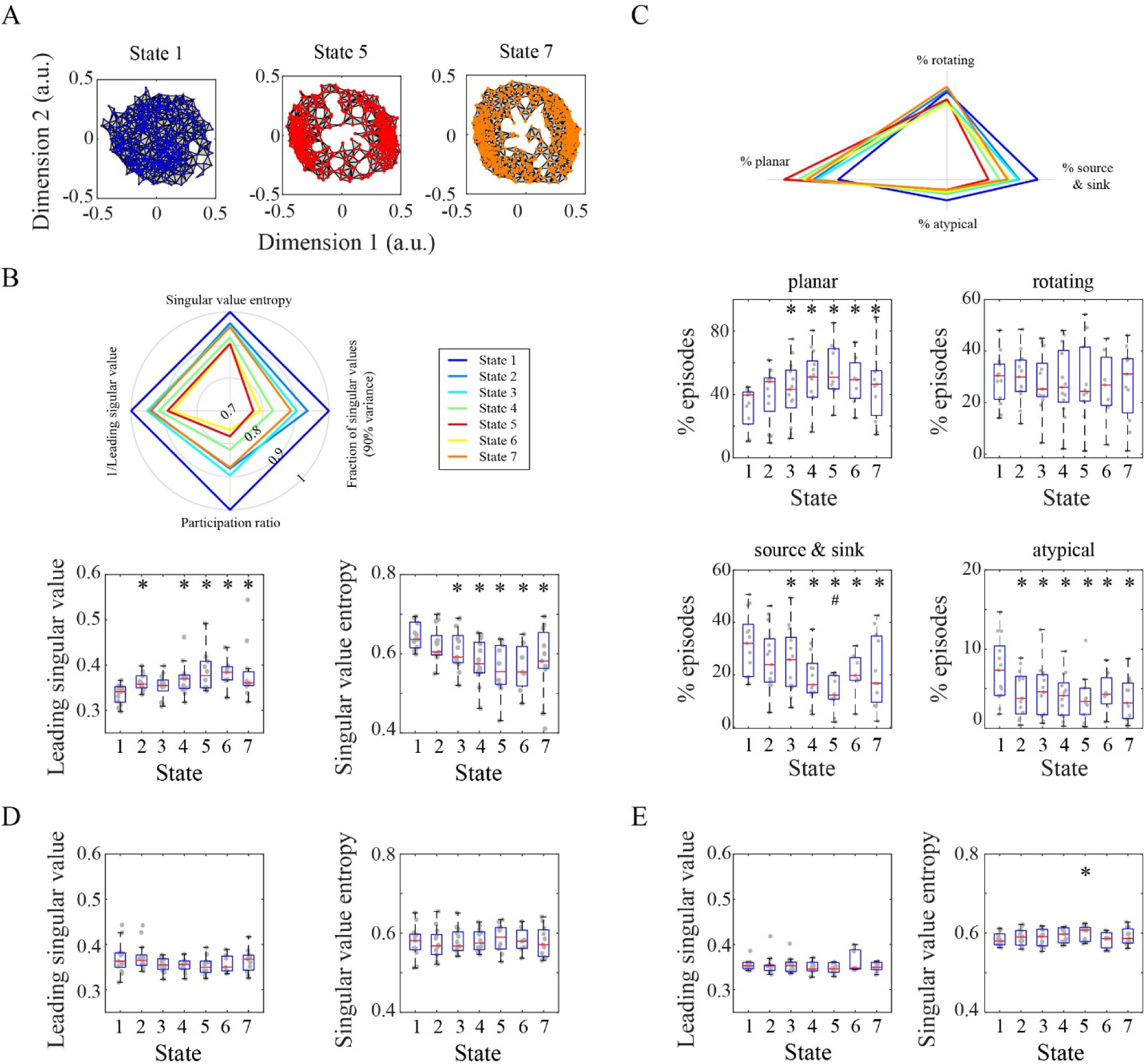
State-dependent reorganization of traveling waves under anesthesia. **A**. Low-dimensional representation of delta-band traveling wave patterns obtained by multidimensional scaling of the episode-by-episode similarity matrix in a representative experiment. Each point represents a delta-wave episode. States 1, 5, and 7 of primary interest are shown. **B**. State-dependent changes in delta-band wave pattern complexity. Box plots and dots show the leading singular value and singular value entropy across experiments. The radar plot summarizes the mean values of these two and two additional complementary measures, with larger values indicating a more diverse repertoire of wave patterns (see Figure S5 for the results of the complementary complexity measures across experiments). **C**. State-dependent changes in the proportion of each delta-band wave type across experiments. Radar plot shows the mean proportion of planar, rotating, source/sink, and atypical wave patterns. **D, E**. State-dependence of theta-(**D**) and gamma-band (**E**) wave pattern complexity. * *p*<0.05/6 vs. State 1. # *p*<0.05/6 vs. State 7, linear mixed-effects models.

Specifically, the largest singular value increased in States 2 and 4-6 (all *p*≤0.006; **Table S2**), whereas singular value entropy decreased in States 3-6 (all *p*≤0.001). Compared with State 5, State 7 showed a partially expanded distribution but remained more constrained than State 1 (**Figure 3A**). Consistent with this observation, both metrics showed a partial reversal relative to State 5, approaching the values observed in States 2 and 3. However, neither metric differed significantly from the intermediate-to-deep anesthetic states (States 4-6; all *p*≥0.224). The largest singular value remained higher (*p*<0.001), and singular value entropy remained lower than in State 1 (*p*<0.001). These findings were corroborated by complementary measures of effective dimensionality (**Figure S5**).

We next examined whether these changes were accompanied by shifts in the composition of wave types. In the awake state, multiple wave types—including planar, rotating, and source/sink (see **Figure S6** for representative examples)—were present in relatively balanced proportions (**Figure 3C**). Under anesthesia, planar waves were dominant, accompanied by a corresponding reduction in more complex wave types, consistent with the increasingly stereotyped and less diverse wave repertoire. The proportion of planar waves increased significantly in States 3-7 (all *p*≤0.001 vs. State 1; **Table S2**), whereas the proportion of source/sink (States 4-7; all *p*≤0.005) and atypical wave patterns (States 2-7; all *p*≤0.004) decreased (**Figure 3D**). The reduction in source/sink wave patterns was partially reversed in State 7 (*p*=0.004 vs. State 5), whereas the proportions of planar and atypical wave patterns did not differ from State 5 after Bonferroni correction (both *p*≥0.008). Comparable analyses of theta- and gamma-band traveling waves revealed little or no state-dependent change in wave-pattern complexity or wave-type composition (**Figure 3D, E** and **Figure S7**).

Together, these results indicate that despite their increased occurrence, delta-band traveling waves under anesthesia are less diverse and more stereotyped, with a greater prevalence of planar propagation patterns. In the paradoxical state, these effects generally show trends toward reversal, although only the anesthetic-induced reduction in source/sink waves is significantly reversed.

### Frequency-dependent organization of planar waves across cortical states

We next examined the directional organization of planar waves across cortical states. Across all planar wave episodes, the distribution of mean propagation directions exhibited a bimodal pattern, with two dominant directions corresponding to feedforward (FF; posterior-to-anterior) and feedback (FB; anterior-to-posterior) propagation across all frequency bands and cortical states (**Figures 4A**, **5A**, and **6A**). We therefore classified planar waves into FF and FB waves for subsequent analyses and quantified the preferred propagation direction (the circular mean across episodes), directional consistency, and relative occurrence of FF and FB waves across cortical states.

**Figure 4.**
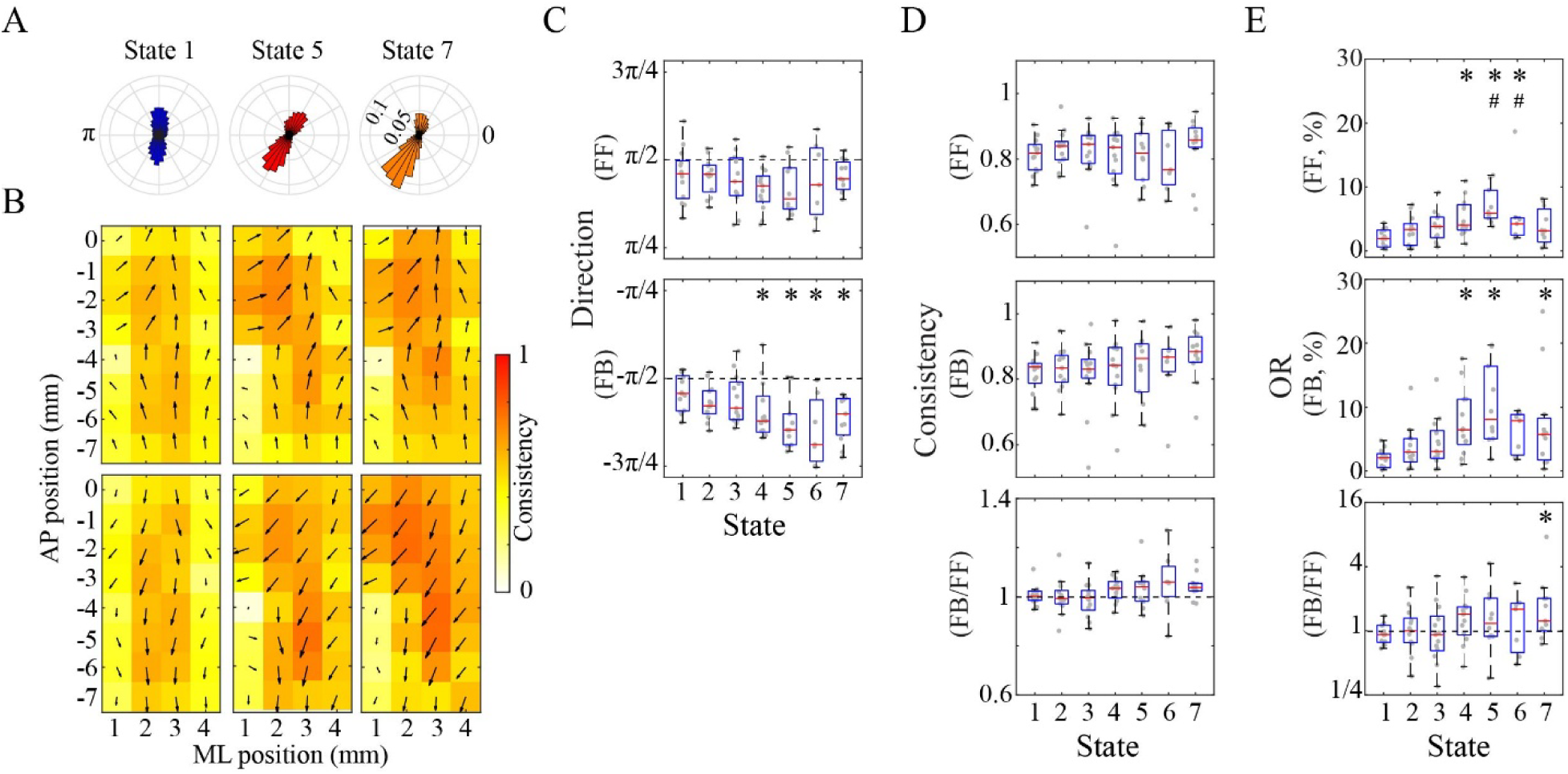
State-dependent modulation of delta-band planar waves by anesthesia. **A**. Distribution of the mean propagation directions of planar wave episodes across electrode locations for three states of primary interest 1, 5, and 7 pooled from all experiments. **B**. Mean feedforward (FF, posterior-to-anterior) and feedback (FB, anterior-to-posterior) planar wave patterns in the three states. Arrows indicate spatial phase-gradient directions across planar wave episodes pooled from all experiments; arrow length and color represent directional consistency (1 - circular variance). **C**. Mean propagation directions of FF and FB waves in seven cortical states. **D**. Between-episode directional consistency of FF and FB waves and their ratio (FB/FF) across states. **E**. Relative occurrence rates (OR) of FF vs. FB planar waves and their ratio (FB/FF) across states. \**p* < 0.05/6 vs. State 1; #*p* < 0.05/6 vs. State 7, linear mixed-effects models. AP, anteroposterior, ML, mediolateral.

**Figure 5.**
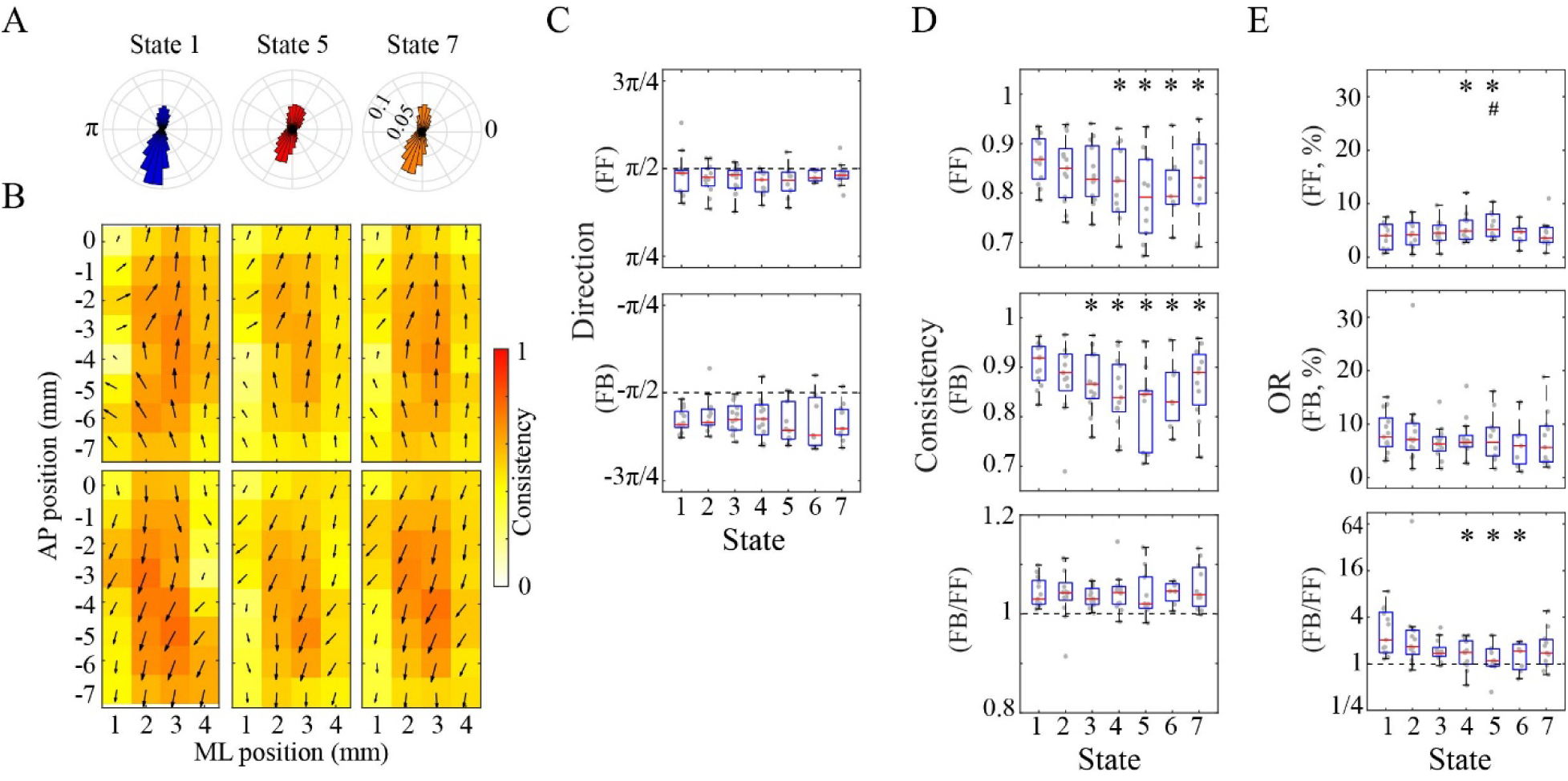
State-dependent modulation of theta-band planar waves by anesthesia. **A**. Distribution of the mean propagation directions of planar wave episodes across electrode locations for three states of primary interest (States 1, 5, and 7) pooled from all experiments. **B**. Mean feedforward (FF, posterior-to-anterior) and feedback (FB, anterior-to-posterior) planar wave patterns in the three states. **C**. Mean propagation directions of FF and FB waves in seven cortical states. **D**. Between-episode directional consistency of FF and FB waves and their ratio (FB/FF) across states. **E**. Relative occurrence rates (OR) of FF vs. FB planar waves and their ratio (FB/FF) across states. In C-E, \**p* < 0.05/6 vs. State 1; #*p* < 0.05/6 vs. State 7, linear mixed-effects models. AP, anteroposterior, ML, mediolateral.

**Figure 6.**
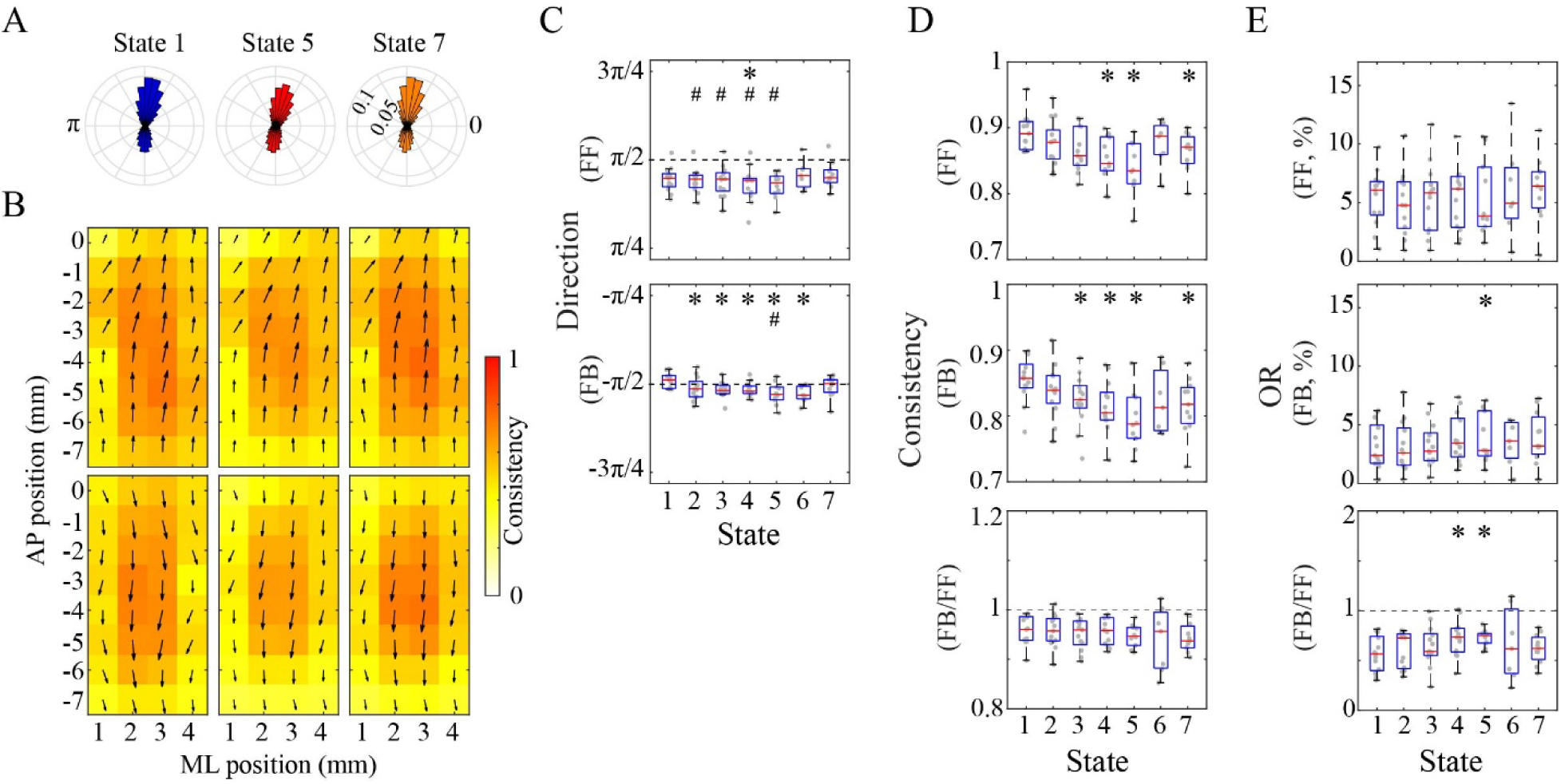
State-dependent modulation of gamma-band planar waves by anesthesia. **A**. Distribution of the mean propagation directions of planar wave episodes across electrode locations for three states of primary interest (States 1, 5, and 7) pooled from all experiments. **B**. Mean feedforward (FF, posterior-to-anterior) and feedback (FB, anterior-to-posterior) planar wave patterns in the three states. **C**. Mean propagation directions of FF and FB waves in seven cortical states. **D**. Between-episode directional consistency of FF and FB waves and their ratio (FB/FF) across states. **E**. Relative occurrence rates (OR) of FF vs. FB waves and their ratio (FB/FF) across states. In C-E, \**p* < 0.05/6 vs. State 1; #*p* < 0.05/6 vs. State 7, linear mixed-effects models. AP, anteroposterior, ML, mediolateral.

### Delta band

In the delta band, the preferred propagation direction of FB waves differed across states, shifting from an anterior-to-posterior direction (−0.54π in the awake state) toward a more diagonally oriented anterior-lateral to posterior-medial direction (−0.69π in State 6) (**Figure 4B, C**). This shift was significant in States 4-7 relative to State 1 (all *p*≤0.006). In State 7, the preferred propagation direction showed a trend toward reversal relative to State 5 (−0.60π), although this difference did not reach statistical significance. FF waves showed a similar but non-significant trend. Despite these shifts in preferred propagation direction, the directional consistency of individual episodes remained similar across cortical states for both FF and FB, indicating stable propagation direction variability across states (**Figure 4D**).

In the awake state, FB and FF waves were balanced in occurrence rate (FB/FF ratio not different from 1, *p*=0.340). Both FF and FB wave occurrence increased significantly (FF: States 4-6; FB: States 4, 5, and 7; all *p*<0.001 vs. State 1) (**Figure 4E**), driven by corresponding increases in both episode frequency and duration (**Figure S8**). In State 7, however, only FF occurrence showed a significant reversal relative to State 5 (*p* < 0.001) (**Figure 4E**), accompanied by corresponding reductions in both episode frequency and duration (**Figure S8**). As a result, the relative balance between FF and FB wave occurrence, which remained similar across States 1-6, shifted significantly toward FB in State 7 compared with State 1 (*p*<0.001; **Figure 4E**).

These results indicate that delta-band planar waves exhibit increased occurrence and shifts in propagation direction in intermediate-to-deep anesthetic states. FF and FB wave occurrence remains relatively balanced across cortical states, except in the paradoxical state, where a reversal of FF wave occurrence shifts the balance toward FB predominance.

### Theta band

In the theta band, both FF and FB waves maintained their awake-state propagation directions (FF: 0.49π, FB: −0.59π) across cortical states (**Figure 5B, C**). In contrast, the directional consistency of individual episodes decreased significantly (FF: States 4-7, *p*<0.001; FB: States 3-7, *p*≤0.005), with a slight, non-significant reversal in State 7 (**Figure 5D**), indicating greater variability around a preserved preferred propagation direction. In addition, FB waves exhibited higher directional consistency than FF waves in the awake state (*p*=0.002 vs. 1), and this difference persisted throughout anesthesia, with no significant effect of state (*p*=0.857) (**Figure 5D**).

In the awake state, FB waves were more prevalent than FF waves (*p*=0.002 vs. 1). While FB occurrence remained unchanged across states, FF wave occurrence increased in States 4-5 (*p*≤0.002 vs. State 1) and reversed in State 7 (*p*=0.599 vs. State 1, *p=*0.008 vs. State 5) (**Figure 5E**). Accordingly, the FB-dominant asymmetry was lost in States 4-6 (*p*≤0.004) and showed a reversal in State 7, which no longer differed significantly from the awake state after Bonferroni correction (*p*=0.028 vs. State 1). A similar pattern was observed for episode frequency, indicating that the changes in FF wave occurrence and FB-dominant occurrence asymmetry were primarily driven by changes in episode frequency rather than episode duration (**Figure S9**).

These results indicate that theta-band planar waves preserve their overall propagation direction despite increased directional variability across individual wave episodes. The FB-dominant asymmetry in wave occurrence present in the awake state is lost in intermediate-to-deep anesthetic states but reversed in the paradoxical state.

### Gamma band

In the gamma band, FF and FB waves propagated predominantly along the anterior-posterior axis in the awake state (FF: 0.45π, FB: −0.49π; **Figure 6B, C**). Small but significant state-dependent shifts in propagation direction were observed (FF: State 4; FB: States 2-6; all *p*≤0.002), and these shifts reversed in State 7 (both *p*≤0.005 vs. State 5). As in the theta band, directional consistency across episodes decreased significantly (FF: States 4, 5 and 7, FB: States 3-5 and 7, all *p*≤0.008), with a slight, non-significant reversal in State 7. In addition, FF waves exhibited higher directional consistency than FB waves in the awake state (*p*=0.002 vs. 1), and this difference persisted throughout anesthesia, with no significant effect of state (*p*=0.329) (**Figure 6D**).

In the awake state, FF waves were more prevalent than FB waves (*p*=0.002 vs. 1) (**Figure 6E**). While FF occurrence remained stable across states, FB wave occurrence showed a mild increase in State 5 (*p*=0.002) but not in State 7 (*p*=0.060 vs State 1). Accordingly, the FF-dominant asymmetry persisted (both *p*≤0.007 vs. 1) but became less pronounced in States 4 and 5 (both *p*≤0.007 vs. State 1) before reversing in State 7, which no longer differed significantly from the awake state (*p*=0.124 vs. State 1). A similar reversal pattern was observed for both episode frequency and duration, with reduced FF-dominant asymmetry in State 5 and re-emergence in State 7 (**Figure S10**).

These results indicate that gamma-band planar waves exhibit small shifts in overall propagation direction and increased directional variability across individual wave episodes. The FF-dominant asymmetry in wave occurrence present in the awake state persists but is attenuated in the deep anesthetic state and is restored in the paradoxical state.

Together, these findings demonstrate that the directional organization of planar waves is frequency dependent, as is its modulation by anesthesia. Distinct from the canonical deep anesthetic state (State 5), the paradoxical state exhibits partial, frequency-dependent reversal of directional organization toward wakefulness (see **Figure 7** for a summary).

**Figure 7.**
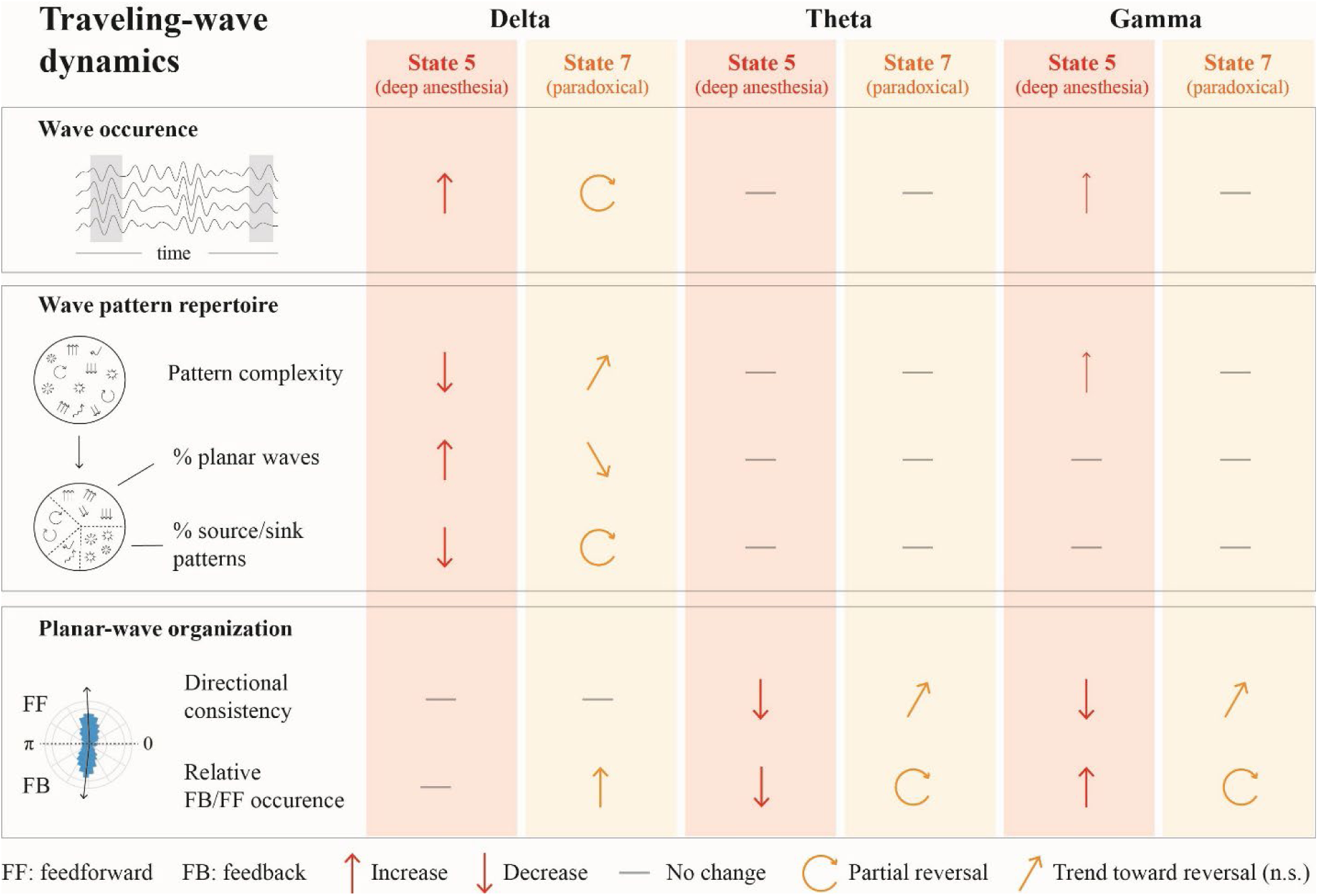
Summary of state- and frequency-dependent modulation of cortical traveling waves by desflurane anesthesia. Changes in traveling-wave dynamics in canonical deep anesthesia (State 5) and the paradoxical state (State 7) are summarized relative to the awake state (State 1) across the delta, theta, and gamma bands. Measures include overall wave occurrence, wave pattern repertoire (pattern complexity and proportions of planar and source/sink patterns), and planar-wave organization (directional consistency and relative occurrence of feedback [FB] vs. feedforward [FF] waves). Upward and downward arrows indicate significant increases and decreases, respectively, relative to State 1; horizontal lines indicate no significant change. Circular arrows indicate partial reversals in State 7 relative to State 5 toward values in the awake state, whereas diagonal arrows indicate similar trends that did not reach statistical significance. n.s., not significant.

## Discussion

Traveling waves reflect the spatial organization and propagation of neural activity across the cortex and are thought to play an important role in coordinating cortical dynamics across space and time. To determine how general anesthesia modulates this spatiotemporal organization, we analyzed traveling waves in hemispheric ECoG recordings at multiple depths of anesthesia with desflurane. Desflurane, a clinically used volatile anesthetic, was chosen because its rapid equilibration allows anesthetic concentration to be systematically controlled across a broad range of anesthetic depths. While an increasing number of studies have examined traveling waves during general anesthesia, most have not systematically investigated their dose or concentration dependence, particularly across anesthetic levels associated with transitions between conscious and unconscious states. Examining traveling waves across graded anesthetic levels may therefore help identify cortical reorganization that accompanies changes in conscious state.

Brain activity is inherently dynamic, spontaneously transitioning among distinct states even in the absence of external stimulation (Deco et al., 2017). General anesthesia alters these ongoing brain-state dynamics, and examining such dynamics may provide further insight into cortical reorganization that is not captured by anesthetic concentration alone. As we have previously shown, cortical states frequently fluctuate even at steady-state anesthetic concentrations (Li and Hudetz, 2026). Building on our prior identification of seven discrete cortical states, here we show that traveling wave dynamics also vary in a state- and frequency-dependent manner.

Compared with normal wakefulness, cortical states predominantly associated with 4-6% desflurane show a selective increase in delta-band wave occurrence, while the repertoire of delta-band patterns becomes more constrained and dominated by stereotyped planar configurations. In contrast, theta- and gamma-band waves show little state-dependent change in overall occurrence or wave pattern complexity but exhibit greater variability in planar-wave propagation direction and frequency-specific alterations in feedback-feedforward organization: theta feedback dominance is lost, whereas gamma feedforward asymmetry persists but becomes less pronounced. These traveling-wave alterations become particularly evident in State 4 (predominantly occurring at 4% desflurane) and persist or become more pronounced in State 5 (predominantly occurring at 6% desflurane). Given that these concentrations fall within or above the range previously associated with loss of righting reflex (Imas et al., 2005b), these changes may reflect cortical reorganization accompanying the transition to behavioral unresponsiveness.

Among the frequency bands examined, delta-band waves show the most prominent anesthesia-induced reorganization. During wakefulness, as in the theta and gamma bands, delta-band waves display a diverse repertoire of planar, rotating, and more localized source/sink patterns, consistent with previous observations that spontaneous cortical activity supports multiple forms of traveling-wave propagation (Roberts et al., 2019; Liang et al., 2023b; Das et al., 2026). In intermediate-to-deep anesthetic states, however, delta-band waves are selectively altered in both their overall occurrence and pattern repertoire: waves are more frequent and longer, while pattern diversity is reduced with a greater predominance of planar waves. These planar waves propagate primarily along opposing directions of the anterior-posterior axis, consistent with prior reports during natural sleep (Massimini et al., 2004) and propofol anesthesia (Murphy et al., 2011), with deep anesthetic state showing a shift toward an anterolateral - posteromedial axis, as observed under isoflurane and urethane anesthesia (Greenberg et al., 2018; Pazienti et al., 2022). The constrained delta-wave repertoire in deep anesthetic state is consistent with prior observations across species and modalities (Dasilva et al., 2021; Bhattacharya et al., 2022; Pazienti et al., 2022; Liang et al., 2023b). Conversely, greater heterogeneity of low-frequency traveling waves has been reported during psychedelic states induced by 5-MeO-DMT (Blackburne et al., 2025). Taken together, these observations suggest that delta-band traveling wave diversity may provide a sensitive measure of brain-state-dependent cortical organization.

Whereas delta-band waves are primarily distinguished by anesthesia-related changes in overall occurrence and pattern repertoire, theta- and gamma-band waves reveal a distinct directional organization of planar waves that is already present during wakefulness. In contrast to the relative balance opposing propagation directions in the delta band, theta- and gamma-band planar waves exhibit frequency-specific directional asymmetry: theta-band waves show feedback (anterior-to-posterior) dominance, whereas gamma-band waves show feedforward (posterior-to-anterior) dominance in occurrence. The directional asymmetry occurs during spontaneous activity, extending previous observations of frequency-dependent traveling wave direction during visually evoked responses (Aggarwal et al., 2022). This organization is consistent with broader evidence linking lower-frequency rhythms to feedback signaling and gamma rhythms to feedforward communication across cortical hierarchies (Michalareas et al., 2016; Richter et al., 2017). The presence of these frequency-specific modes of wave propagation in the absence of external input suggests that they reflect the intrinsic organization of cortical activity.

General anesthesia alters this intrinsic directional organization in a frequency-dependent manner. In intermediate-to-deep anesthetic states, theta-band feedback dominance is lost, whereas gamma-band feedforward asymmetry persists, although it becomes less pronounced. This dissociation is consistent with prior work showing that isoflurane selectively impairs low-frequency feedback while preserving high-frequency feedforward connectivity in the fly brain (Cohen et al., 2018). More broadly, these findings align with evidence that anesthetics preferentially disrupt feedback or recurrent connectivity across cortex (Imas et al., 2005a; Ku et al., 2011; Boly et al., 2012; Lee et al., 2013; Murphy et al., 2019). Despite these frequency-specific changes in directional asymmetry, theta- and gamma-band planar waves share a reduction in directional consistency, indicating less stable propagation directions under anesthesia. This increased directional variability is absent in delta-band planar waves, which instead show a more constrained overall pattern repertoire. Together, these findings indicate that anesthesia reorganizes distinct aspects of traveling wave dynamics in a frequency-dependent manner.

Against the frequency-dependent reorganization observed in canonical deep anesthetic state, the paradoxical state is particularly notable because traveling wave properties generally shift toward those observed during lighter anesthesia or wakefulness, despite occurring predominantly at the highest desflurane concentration examined here. In particular, although the overall delta-wave pattern repertoire shifts toward lighter anesthetic states without reaching statistical significance, the prevalence of source/sink patterns shows a significant reversal. The partial restoration of theta feedback and gamma feedforward asymmetries further suggests that aspects of the frequency-specific directional organization observed during wakefulness are re-established in the paradoxical state. However, whether this reorganization reflects any restoration of conscious processing or represents a distinct form of cortical activation without true awakening remains to be determined (Destexhe et al., 2007; Li and Hudetz, 2025). Together with our previous findings of reduced delta power and elevated spatiotemporal Lempel-Ziv complexity (Li and Hudetz, 2026), these results further support the paradoxical state as a distinct dynamical regime within deep anesthesia. Traveling-wave dynamics extend these previous findings by providing complementary information about the spatiotemporal organization of this state beyond that captured by spectral power or complexity alone.

The present study has several methodological limitations. First, the traveling wave analyses are based on phase propagation patterns derived from spontaneous recordings and therefore do not directly establish causal interactions or directional information flow. Future studies incorporating directed connectivity or perturbational approaches will be necessary to clarify the relationship between traveling wave dynamics and underlying causal interactions in cortical networks.

Second, the spatial coverage of our hemispheric electrocorticography recordings is limited, constraining our ability to fully characterize large-scale cortical wave propagation, particularly spatially extended patterns such as rotating waves, and potentially underestimating the diversity of wave patterns. Third, hemispheric ECoG recordings capture large-scale cortical dynamics but provide limited information about local network activity. Multi-scale studies combining ECoG with local field potential or unit recordings within the same subjects (Buzsaki et al., 2012; Yue et al., 2019) will be important for clarifying the neuronal mechanisms underlying traveling wave dynamics. Fourth, although anesthetic modulation of traveling waves is robust across frequency bands, cross-frequency co-occurrence of traveling wave episodes is weak and variable in the present dataset. This may in part reflect our focus on spontaneous activity, in which cross-frequency coordination of traveling waves may be less pronounced than under stimulus-driven conditions. Finally, this study focuses on desflurane anesthesia in rats; future work is needed to determine whether the observed traveling wave dynamics generalize across anesthetic agents with different pharmacological mechanisms and across species.

In summary, desflurane anesthesia differentially modulates cortical traveling waves across frequency bands, with the canonical intermediate-to-deep anesthetic states characterized by more constrained delta-wave patterns and altered feedback-feedforward organization of theta- and gamma-band waves. These changes are partially reversed in the paradoxical state despite its predominant occurrence at the highest anesthetic concentration studied here, suggesting a shift toward aspects of the traveling wave organization observed during wakefulness. These findings demonstrate the usefulness of traveling waves for studying how large-scale cortical dynamics reorganize across brain states under anesthesia and provide new insight into spatiotemporal dynamics relevant to anesthetic modulation of consciousness.

## Acknowledgements

Research reported in this publication was supported by the National Institute of General Medical Sciences of the National Institutes of Health under award number R01-GM056398 and the Center for Consciousness Science, Department of Anesthesiology, University of Michigan Medical School, Ann Arbor, Michigan, USA. The content is solely the responsibility of the authors and does not necessarily represent the official views of the National Institutes of Health. The authors express their gratitude to Dr. Shiyong Wang for his assistance in performing the experiments, to Drs. Gábor Juhász and Zsolt Borhegyi, Eötvös Loránd University, Budapest, Hungary and to Dr. Zoltán Fekete, Pázmány Péter Catholic University, Budapest for their advice for the use of flexible polymer electrodes.

## References

Aggarwal A, Luo J, Chung H, Contreras D, Kelz MB, Proekt A (2024) Neural assemblies coordinated by cortical waves are associated with waking and hallucinatory brain states. Cell Rep 43:114017.

Aggarwal A, Brennan C, Luo J, Chung H, Contreras D, Kelz MB, Proekt A (2022) Visual evoked feedforward-feedback traveling waves organize neural activity across the cortical hierarchy in mice. Nat Commun 13:4754.

Balasubramanian K, Arce-McShane FI, Dekleva BM, Collinger JL, Hatsopoulos NG (2023) Propagating motor cortical patterns of excitability are ubiquitous across human and non-human primate movement initiation. iScience 26:106518.

Bardon AG, Ballesteros JJ, Brincat SL, Roy JE, Mahnke MK, Ishizawa Y, Brown EN, Miller EK (2025) Convergent effects of different anesthetics on changes in phase alignment of cortical oscillations. Cell Rep 44:115685.

Berens P (2009) CircStat: A MATLAB Toolbox for Circular Statistics. J Stat Softw 31:1–21.

Bhattacharya S, Donoghue JA, Mahnke M, Brincat SL, Brown EN, Miller EK (2022) Propofol Anesthesia Alters Cortical Traveling Waves. J Cogn Neurosci 34:1274–1286.

Blackburne G, McAlpine RG, Fabus M, Liardi A, Kamboj SK, Mediano PAM, Skipper JI (2025) Complex slow waves in the human brain under 5-MeO-DMT. Cell Rep 44:116040.

Boly M, Moran R, Murphy M, Boveroux P, Bruno MA, Noirhomme Q, Ledoux D, Bonhomme V, Brichant JF, Tononi G, Laureys S, Friston K (2012) Connectivity changes underlying spectral EEG changes during propofol-induced loss of consciousness. J Neurosci 32:7082–7090.

Buzsaki G, Anastassiou CA, Koch C (2012) The origin of extracellular fields and currents - EEG, ECoG, LFP and spikes. Nature Reviews Neuroscience 13:407–420.

Cohen D, van Swinderen B, Tsuchiya N (2018) Isoflurane Impairs Low-Frequency Feedback but Leaves High-Frequency Feedforward Connectivity Intact in the Fly Brain. eNeuro 5.

Cruddas J, Pang JC, Fornito A (2026) Cortical traveling waves in time and space: Physics, physiology, and psychology. Neuron 114:985–1005.

Das A, Zabeh E, Ermentrout B, Jacobs J (2026) Planar, spiral, and concentric traveling waves distinguish behavioral states in human memory. Nat Commun 17:5143.

Dasilva M, Camassa A, Navarro-Guzman A, Pazienti A, Perez-Mendez L, Zamora-Lopez G, Mattia M, Sanchez-Vives MV (2021) Modulation of cortical slow oscillations and complexity across anesthesia levels. Neuroimage 224:117415.

Davis ZW, Muller L, Martinez-Trujillo J, Sejnowski T, Reynolds JH (2020) Spontaneous travelling cortical waves gate perception in behaving primates. Nature 587:432–436.

Deco G, Kringelbach ML, Jirsa VK, Ritter P (2017) The dynamics of resting fluctuations in the brain: metastability and its dynamical cortical core. Scientific Reports 7.

Destexhe A, Hughes SW, Rudolph M, Crunelli V (2007) Are corticothalamic ‘up’ states fragments of wakefulness? Trends Neurosci 30:334–342.

Fedor FZ, Zatonyi A, Cserpan D, Somogyvari Z, Borhegyi Z, Juhasz G, Fekete Z (2020) Application of a flexible polymer microECoG array to map functional coherence in schizophrenia model. MethodsX 7:101117.

Franks NP (2008) General anaesthesia: from molecular targets to neuronal pathways of sleep and arousal. Nat Rev Neurosci 9:370–386.

Greenberg A, Abadchi JK, Dickson CT, Mohajerani MH (2018) New waves: Rhythmic electrical field stimulation systematically alters spontaneous slow dynamics across mouse neocortex. Neuroimage 174:328–339.

Hudetz AG, Mashour GA (2016) Disconnecting Consciousness: Is There a Common Anesthetic End Point? Anesth Analg 123:1228–1240.

Hudson AE, Calderon DP, Pfaff DW, Proekt A (2014) Recovery of consciousness is mediated by a network of discrete metastable activity states. Proceedings of the National Academy of Sciences 111:9283.

Imas OA, Ropella KM, Ward BD, Wood JD, Hudetz AG (2005a) Volatile anesthetics disrupt frontal-posterior recurrent information transfer at gamma frequencies in rat. Neuroscience Letters 387:145–150.

Imas OA, Ropella KM, Ward BD, Wood JD, Hudetz AG (2005b) Volatile anesthetics enhance flash-induced gamma oscillations in rat visual cortex. Anesthesiology 102:937–947.

Ito J, Nikolaev AR, van Leeuwen C (2007) Dynamics of spontaneous transitions between global brain states. Human Brain Mapping 28:904–913.

Ku SW, Lee U, Noh GJ, Jun IG, Mashour GA (2011) Preferential Inhibition of Frontal-to-Parietal Feedback Connectivity Is a Neurophysiologic Correlate of General Anesthesia in Surgical Patients. Plos One 6.

Lee H, Wang SY, Hudetz AG (2020) State-Dependent Cortical Unit Activity Reflects Dynamic Brain State Transitions in Anesthesia. Journal of Neuroscience 40:9440–9454.

Lee U, Ku S, Noh G, Baek S, Choi B, Mashour GA (2013) Disruption of Frontal–Parietal Communication by Ketamine, Propofol, and Sevoflurane. Anesthesiology 118:1264–1275.

Li D, Hudetz AG (2025) Anesthesia alters complexity of spontaneous and stimulus-related neuronal firing patterns in rat visual cortex. Neuroscience 565:440–456.

Li D, Hudetz AG (2026) Dynamic Electrocortical States and Paradoxical Complexity during Desflurane Anesthesia. Anesthesiology 145:373–386.

Li D, Vlisides PE, Kelz MB, Avidan MS, Mashour GA, Group ftRS (2019) Dynamic Cortical Connectivity during General Anesthesia in Healthy Volunteers. Anesthesiology: The Journal of the American Society of Anesthesiologists 130:870–884.

Liang W, Balasubramanian K, Papadourakis V, Hatsopoulos NG (2023a) Propagating spatiotemporal activity patterns across macaque motor cortex carry kinematic information. Proc Natl Acad Sci U S A 120:e2212227120.

Liang Y, Liang J, Song C, Liu M, Knopfel T, Gong P, Zhou C (2023b) Complexity of cortical wave patterns of the wake mouse cortex. Nat Commun 14:1434.

Liang YQ, Song CC, Liu MX, Gong PL, Zhou CS, Knöpfel T (2021) Cortex-Wide Dynamics of Intrinsic Electrical Activities: Propagating Waves and Their Interactions. Journal of Neuroscience 41:3665–3678.

Massimini M, Huber R, Ferrarelli F, Hill S, Tononi G (2004) The sleep slow oscillation as a traveling wave. Journal of Neuroscience 24:6862–6870.

Michalareas G, Vezoli J, van Pelt S, Schoffelen JM, Kennedy H, Fries P (2016) Alpha-Beta and Gamma Rhythms Subserve Feedback and Feedforward Influences among Human Visual Cortical Areas. Neuron 89:384–397.

Muller L, Chavane F, Reynolds J, Sejnowski TJ (2018) Cortical travelling waves: mechanisms and computational principles. Nat Rev Neurosci 19:255–268.

Muller L, Busch AN, Davis ZW, Reynolds JH (2026) Neural traveling waves in cortex: Network mechanisms and potential roles in neural computation. Neuron.

Muller L, Piantoni G, Koller D, Cash SS, Halgren E, Sejnowski TJ (2016) Rotating waves during human sleep spindles organize global patterns of activity that repeat precisely through the night. Elife 5.

Murphy C, Krause B, Banks M (2019) Selective effects of isoflurane on cortico-cortical feedback afferent responses in murine non-primary neocortex. Br J Anaesth 123:488–496.

Murphy M, Bruno MA, Riedner BA, Boveroux P, Noirhomme Q, Landsness EC, Brichant JF, Phillips C, Massimini M, Laureys S, Tononi G, Boly M (2011) Propofol anesthesia and sleep: a high-density EEG study. Sleep 34:283–291A.

National Research Council (2011) Guide for the Care and Use of Laboratory Animals, 8 Edition. Washington, DC: National Academies Press.

Pazienti A, Galluzzi A, Dasilva M, Sanchez-Vives MV, Mattia M (2022) Slow waves form expanding, memory-rich mesostates steered by local excitability in fading anesthesia. iScience 25:103918.

Pazienti A, Müller M, Bosman CA, Olcese U, Mattia M (2025) Circuit-level dynamics and propagation of slow wave activity modulate their interplay during the awakening process. Iscience 28.

Percie du Sert N et al. (2020) Reporting animal research: Explanation and elaboration for the ARRIVE guidelines 2.0. PLoS Biol 18:e3000411.

Purdon PL, Sampson A, Pavone KJ, Brown EN (2015) Clinical Electroencephalography for Anesthesiologists: Part I: Background and Basic Signatures. Anesthesiology 123:937–960.

Richter CG, Thompson WH, Bosman CA, Fries P (2017) Top-Down Beta Enhances Bottom-Up Gamma. J Neurosci 37:6698–6711.

Roberts JA, Gollo LL, Abeysuriya RG, Roberts G, Mitchell PB, Woolrich MW, Breakspear M (2019) Metastable brain waves. Nat Commun 10.

Rubner Y, Tomasi C, Guibas LJ (2000) The Earth Mover’s Distance as a metric for image retrieval. Int J Comput Vision 40:99–121.

Townsend RG, Solomon SS, Chen SC, Pietersen AN, Martin PR, Solomon SG, Gong P (2015) Emergence of complex wave patterns in primate cerebral cortex. J Neurosci 35:4657–4662.

Xu YB, McInnes A, Kao CH, D’Rozario A, Feng JF, Gong PL (2025) Spatiotemporal dynamics of sleep spindles form spiral waves that predict overnight memory consolidation and age-related memory decline. Communications Biology 8.

Yilmaz U (2009) The Earth Mover’s Distance. In: MATLAB Central File Exchange.

Yue L, Zhang F, Lu X, Wan Y, Hu L (2019) Simultaneous Recordings of Cortical Local Field Potentials and Electrocorticograms in Response to Nociceptive Laser Stimuli from Freely Moving Rats. J Vis Exp.

Zarr VM, Davis TS, House PA, Greger B, Davis ZW, Smith EH (2026) Propofol-induced loss of responsiveness reorganizes cortical traveling waves in the human brain. bioRxiv.

